# Neurocognitive characteristics of perceptual feedback processing differ in autism

**DOI:** 10.1101/2024.07.23.604831

**Authors:** Nathaniel J. Zuk, Jaroslav Y. Slobodskoy-Plusnin, Yarden Weiss, VishnuPriya Sampath, Kira M. Düsterwald, Athena Akrami, Merav Ahissar

## Abstract

Autistic individuals often show reduced adaptation to changing environmental demands, but the neurocognitive mechanisms underlying this difference remain unclear. We asked whether it may be explained by reduced sensitivity to external feedback and changes in task difficulty. Autistic and non-autistic adults of either sex, matched for cognitive ability, performed a pitch discrimination task in which their response was immediately followed by feedback. Accuracy was similar between the two groups, yet autistic participants exhibited a markedly reduced feedback-related negativity (FRN), an event-related EEG potential differentiating explicit negative versus positive feedback. They also showed weaker modulation of feedback-evoked EEG responses by perceptual difficulty. Lastly, they showed reduced trial-by-trial behavioral updating of perceptual bias following recent correct and easy trials. These effects were replicated in a follow-up experiment conducted 1–2 years later. When combined they classified autistic and non-autistic participants with about 80% accuracy. Together, our findings indicate that multiple processes for evaluating perceptual decisions and adjusting subsequent behavior differ in autism and likely contribute to slow updating in changing contexts.

**Significance Statement:** Autism is associated with reduced adaptation to changing environments in both behavior and neural signals, but the underlying neurocognitive mechanisms remain unclear. Here, we show that autistic individuals exhibit a reduced feedback-related negativity (FRN), an EEG component that is present when receiving negative instead of positive feedback. They also show weaker neural modulation by perceptual difficulty and reduced trial-by-trial behavioral updating of perceptual bias following recent correct and easy trials. These effects were replicated in a follow-up study. When combined, they classified autistic and non-autistic participants with high accuracy. These findings suggest that reduced flexibility in autism is associated with reduced processing of external feedback and task difficulty.

## Introduction

Individuals with autism spectrum disorder (ASD) often show atypical adaptation to changing environmental demands (Sinha et al., 2014; Van de Cruys et al., 2014; Lieder et al., 2019). In perceptual discrimination tasks, observers implicitly form beliefs about the statistical structure of experienced stimuli and make predictions regarding upcoming inputs (Ashourian and Loewenstein, 2011). These predictions are updated continuously on a trial-by-trial basis (Akrami et al., 2018; Lieder et al., 2019). Efficient adaptation, however, requires not only updating perceptual predictions, but also evaluating the reliability of those predictions and the information used to update them. Several studies suggest that these updating processes are altered in ASD (Lieder et al., 2019; Sapey-Triomphe et al., 2023; Noel et al., 2025; but see Bosch et al., 2022). Related differences have also been observed in tasks requiring continuous performance monitoring and behavioral adjustment following errors (South et al., 2010, 2012; D’Cruz et al., 2016; Doenyas et al., 2019; Vishne et al., 2021; Kasten et al., 2023). Together, these findings suggest that atypical incorporation of ongoing information into internal representations may contribute to altered beliefs, and therefore maladaptive behavior, in ASD.

One process that may contribute to altered belief updating in ASD is evaluation of feedback (Noel et al., 2021). Trial-by-trial feedback provides explicit information regarding response correctness and can influence subsequent perceptual decisions. A substantial literature has examined neural markers of performance monitoring in this context with EEG. When participants are implicitly aware of their errors without any external feedback—for example, participants detect that they have made an error in speed-dependent motor tasks, as in the flanker task (Gehring et al., 1993, 1995, 2018)— the error-related negativity is elicited shortly after the response. A related EEG component, the feedback-related negativity (FRN), is elicited by explicit negative or worse-than-expected feedback (Miltner et al., 1997). Several studies report reduced error-related negativity amplitudes in ASD (Vlamings et al., 2008; Sokhadze et al., 2010; South et al., 2010; Hüpen et al., 2016) and a reduced FRN in ASD (Bellebaum et al., 2014), although this finding is not universal (Henderson et al., 2006; Groen et al., 2008; Larson et al., 2011; for review see Hüpen et al., 2016).

The lack of agreement between ASD studies might hinge on the utility of feedback during the task and its relevance in the specific protocol. In non-autistics, perceptual priors are weighted by one’s confidence in their utility for making judgements (Samaha et al., 2019) which would be affected by additional knowledge, provided by feedback, of whether one’s choices are correct or not. Continuously weighting perceptual priors based on one’s confidence is also important for recognizing changes in task contexts (Heald et al., 2021). Importantly, the same brain areas involved in generating the error-related negativity and FRN are also implicated in representing trial history, outcomes, and predictability, as well as updating subsequent actions (Debener et al., 2005; Behrens et al., 2007; Beckmann et al., 2009; Stevens et al., 2011; Sapey-Triomphe et al., 2023; Oesch et al., 2026). The impact of feedback on choice confidence, and consequently updating priors, may both differ in ASD, yet whether these co-occur in a single task has been unexplored. Given previous work showing cognitive differences of perceptual updating between ASD and non-autistic participants in a discrimination task (Lieder et al, 2019), we reasoned that such a task would allow us to directly evaluate group differences in feedback processing and its relation to the use of perceptual priors.

Using this serial pitch discrimination task, we measured the FRN, neural modulation by perceptual difficulty, and behavioral updating based on recent perceptual history. ASD participants showed markedly reduced FRN responses, alongside weaker modulation by perceptual difficulty and recent perceptual history. These effects were replicated in a follow-up study conducted 1–2 years later. Our results indicate that autistic individuals show reduced propensity for processing external and internal feedback, which affects updating priors in a perceptual task.

## Materials and Methods

All experimental methods were approved by the European Research Council and the Ethics Committee of the Hebrew University.

### Study 1

#### Participants

35 non-autistic and 34 ASD individuals participated in this experiment. Participants with autism (and no language difficulty, as assessed by self-report and current reading scores) were recruited through clinicians, support centers, and designated facilities. All had community-based diagnoses, conducted by multi-disciplinary teams of medical specialists who had used various standard diagnostic tools, including ADI-R (Lord et al., 1994), ADOS-G (Lord et al., 2000), and CARS (Schopler et al., 1980) to ensure that all participants met DSM-IV criteria (American Psychiatric Association, 2000). One ASD participant was rejected from further analyses because of poor performance which suggested that they did not understand the task, leaving 33 ASD included in our analyses. Except for one participant in each group, all participants had at most four years of musical training, so that differences in prior musical education would not affect performance in pitch discrimination. The 35 non-autistics and 33 ASD participants that were included in subsequent analyses were also matched for age, gender (57 males and 11 females total), reasoning, and linguistic skills. Reasoning was measured with Block Design, a standardized test of visuospatial reasoning (Wechsler, 2008) (see Table 1). Basic linguistic skills were measured by accuracy of reading a list of 24 pseudo-words in Hebrew (Deutsch and Bentin, 1996). Participants were instructed to read the words aloud, as quickly and accurately as possible.

**Table 1:** Demographic information for the participants of the Study 1. For age, Block Design, and AQ50 the mean is specified for each group, with the standard deviation in parentheses. The age range is also included in brackets for each group. ** p < 0.01 based on a Student’s t-test.

By contrast, and as expected, there was a significant difference in the scores of the self-reported autism questionnaire, AQ50 (Autism Quotient questionnaire; Baron-Cohen et al., 2001), with an effect size of Cohen’s d = 1.10.

#### Experiment design

Experiments were coded using PsychToolbox in Matlab (Mathworks). In each trial, participants were sequentially presented with two tones 50 ms in duration with 5 ms onset and offset sine ramps. Tone frequencies ranged from 800 to 1250 Hz and were sampled so that both tones in a pair were either above or below 1000 Hz (as in Jaffe-Dax et al., 2017). The ratio of the two tone frequencies ranged from 1.01 to 1.25 (0.17 to 3.86 semitones). Tone onsets were separated by 2 seconds, resulting in a working memory delay (inter-stimulus-interval) of 1.95 seconds. After the second tone, participants had to specify which tone had a higher pitch. Upon response, participants were immediately presented with visual feedback for 400 ms: “Correct” in green if they got the correct answer, “Incorrect” in red if they were wrong, or “Miss” in red. The feedback was presented on a 24-inch monitor at a viewing distance of approximately 60 cm, resulting in a text size of approximately 1° of visual angle. Feedback words were displayed with a text height of 1 cm, ensuring clear readability without disproportionately enhancing visual salience. Participants had 1300 ms to respond after the onset of the second tone. If they did not respond within this interval, then they saw “Miss” in red on screen for 400 ms. This “time out” interval was designed to avoid response pressure: the median response time across all participants was 646 ms with a range of 398 ms to 923 ms, well before the 1300 ms cut-off. The next trial was always presented four seconds after the first tone onset of the previous trial (see Figure 1A), irrespective of the speed of the participant’s response. The inter-trial interval (2 s after the second tone) was set to be equal to the within-trial inter-stimulus interval (2 s) in order to reduce the effect of within-trial adaptation, which might differ between the groups as it relates to implicit learning (Jaffe-Dax et al., 2017). The sounds were presented over Sennheiser HD 25 headphones. The volume was adjusted to a comfortable sound level for each participant at the start of the experiment.

**Figure 1:**
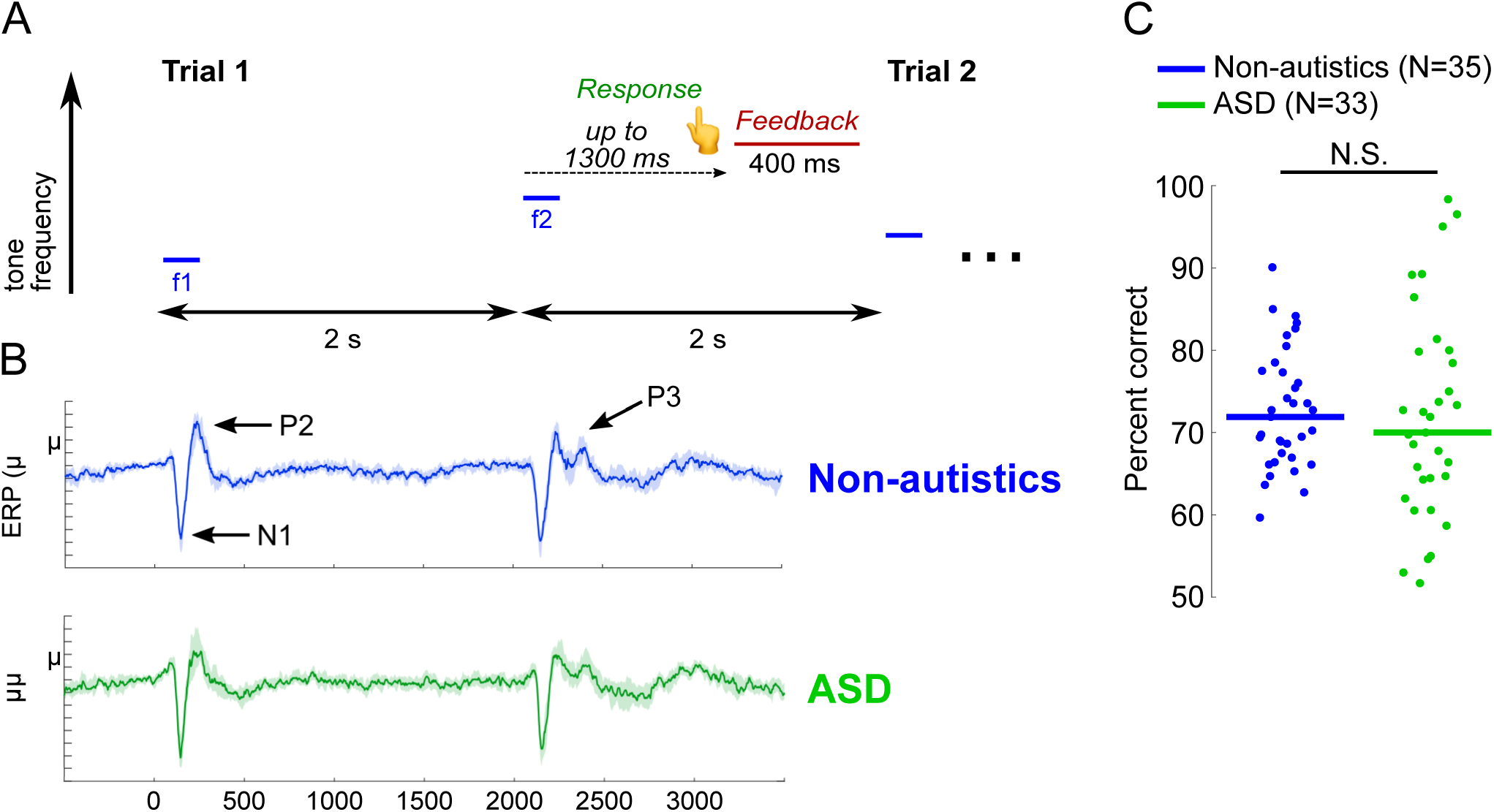
Non-autistics and ASD show similar performance and similar event-related potentials (ERP) to the tones in each trial. (A) A schematic diagram of a trial’s temporal structure. Participants performed a two-tone discrimination task in which they had to determine if the pitch of the second tone was higher than that of the first. Visual feedback ("Correct" or "Incorrect") was given immediately upon response in every trial. (B) Grand average response (median of participants’ median ERPs, see Methods) for electrode Cz. Time zero denotes the onset of the first tone on each trial. Thick lines show the median across participants’ Huber M-estimator ERPs. Shaded regions show the 95% confidence interval of the group median using a bootstrap resampling across participants. Non-autistics are plotted in blue (top) and ASD in green (bottom). Dashed vertical lines mark the onset of each tone. The three main response components, N1, P2 and P3, are marked on the top plot. (C) Distributions of participants’ performance accuracy, in each group. Each dot denotes the percent correct of an individual participant, the bars indicate the median in each group. Median performance was similar in the two groups (no significant difference in a rank-sum test).

Before starting the formal experiment, participants completed a training set of 10-30 trials, presented in blocks of 10 trials with a break in between. To make the task easier during these training trials, the frequency range of the tones was increased to 600 – 1600 Hz, and the tone ratios were randomly selected between 1.35 – 1.4 (equivalent to 5.20 – 5.83 semitones). Participants proceeded to the main experiment upon reaching 8/10 trials correct in a training block, or upon completing the three 10-trial training blocks even if this threshold performance was not attained (this was the case for 7/33 ASD participants and 5/35 non-autistics). In the main experiment, participants performed 121 trials without breaks.

#### Behavioral analysis

Recent trials biased the decision of participants. This recency bias can be viewed as a contraction of the perceptual memory of the first tone in the trial (whose representation is noisier by the time the second tone is presented) towards the expected frequency, namely the geometric mean of the tone frequencies presented in the previous trial. Consequently, in trials in which the first tone’s frequency is closer than the second tone’s frequency to the geometric mean of the previous trial, performance improves, as the mental representation of the first tone becomes more distant from that of the second tone in the trial (termed “bias+” trials). By contrast, in trials in which the first tone is farther from the previous trial’s frequencies, the representation of the distance between the tones decreases, hampering performance (“bias-” trials). Previous work has shown that people with ASD are less biased by the recent trial (Lieder et al., 2019).

Performance was assessed by calculating the proportion of correct trials, ignoring misses. To quantify the effect of recent history on choice bias, we split trials into “bias+” and “bias-” trials, and we calculated the difference in performance between these sets of trials:

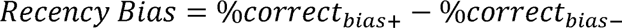

To understand the effects of external feedback (“Correct” or “Incorrect” trials) and trial difficulty on subsequent behavior, we further grouped trials based on the participant’s accuracy (Correct/Incorrect) and the trial difficulty (Easy/Hard trials). For trial difficulty, easy versus hard trials were identified with respect to the study median frequency difference which was 1.242 semitones. Trials with a smaller frequency difference were tagged “hard” whereas trials with larger frequency difference were tagged “easy”. We asked how implicit strategies differed between trials that followed easy ones and trials that followed difficult ones.

Because the recency bias analysis depended on both previous trial difficulty and previous trial accuracy, the number of trials used in this analysis varied across participants. For non-autistics the median number of trials was 31 conditioned on the previous trial being hard and correct (range of 13 – 44 trials) and the median was 36 trials when previously easy and correct (24 – 53). For ASD participants, the median was 27 trials when previously hard and correct (15 – 54) and the median was 33 trials when previously easy and correct (10 – 50).

#### EEG recording and preprocessing

Biosemi ActiveTwo with a 32-channel layout was used to record EEG while participants performed the task. Additionally, external channels were recorded bilaterally on the mastoids, to the outside of each eye, above each eye, and on the nose (7 external electrodes total). The EEG data was digitized at 512 Hz.

All preprocessing was done in Matlab. After recording, the EEG data was filtered using a 4^th^ order zero-phase Butterworth filter with -3 dB cutoff frequencies of 1-30 Hz. Eyeblinks and eye motion artifacts were isolated using fastICA (Hyvärinen and Oja, 2000) applied to the 32 EEG channels as well as the externals and then removed. The data was re-referenced to the average of the mastoid channels. If one of the mastoid channel’s variance exceeded the median plus two times the interquartile range of the main 32 EEG channels (not including the externals), the mastoid channel was deemed too noisy and only the other mastoid channel was used for reference. In two non-autistic participants, both mastoid channels exceeded this range, and we used the average of neighboring P7 and P8 electrodes instead.

#### Calculating evoked responses

All analyses of the EEG were done using channel Cz referenced to the mastoids. Event-related potentials (ERP) to the stimuli and the button presses (the participant’s response in the task) were first calculated individually for each participant. ERPs to the stimuli were baselined to the interval of -100 to 0 ms before the stimulus, and ERPs to the button presses (and feedback presentation) were baselined -400 to -200 ms; we chose a different interval before the button press to avoid a negativity that appeared between -200 to 0 ms in some participants.

To reduce the influence of extreme single-trial EEG amplitudes while retaining all available trials, ERPs were estimated using the Huber M-estimator across trials, separately for each participant, condition, and channel. The Huber M-estimator is a robust estimator of central tendency: values close to the estimated center are treated similarly to conventional averaging, whereas observations with large residuals are down-weighted (Huber, 1964; McWilliams et al., 2026). Thus, the procedure does not discard trials but reduces the influence of occasional extreme amplitudes. No temporal smoothing was applied to the individual ERP waveforms used for statistical analyses. We also verified that the main ERP results were robust to alternative ERP estimators, including conventional mean ERPs, pointwise median ERPs, 20% total trimmed means, 20% total Winsorized means (Rousselet et al., 2017), with and without temporal smoothing.

For plotting group-level ERPs, the participant-level ERP waveforms were summarized using the median across participants. Shaded regions in the ERP plots (e.g. Figure 1B) indicate bootstrapped 95% confidence intervals of the group median, computed by resampling participants with replacement 1000 times. For statistical analyses of ERP magnitudes, minima and maxima were extracted from the individual participant-level Huber-estimated ERPs.

Because our individual EEG analyses were conditioned on trial-specific (hard versus easy trial) and behavioral (correct versus incorrect trial) factors, the number of trials that were in each analysis varied across participants but were comparable between groups. For the feedback-related negativity (FRN) analysis, non-autistics had a median of 86 correct trials (range of 66 – 108) and a median of 33 incorrect trials (12 - 47), while ASD participants had a median of 79 correct trials (31 – 107) and a median of 35 incorrect trials (9 – 55). For the peak-to-peak analysis of the responses to feedback, non-autistics had a median of 57 hard trials (48 – 75) and a median of 60 easy trials (45 – 69). ASD participants had a median of 57 hard trials (25 – 73) and a median of 57 easy trials (20 – 69).

As an additional distributional robustness analysis, we performed shift-function and difference-asymmetry analyses following Rousselet et al. (2017). Shift functions were used to compare non-autistic and ASD groups across quantiles of the individual-measure distributions, rather than only at the group mean or median. Difference-asymmetry functions were used to assess whether the distribution of all pairwise differences between non-autistic and ASD magnitudes reflected a broad shift or was driven by asymmetric tails. These analyses were applied to the three primary measures: FRN magnitude, hard-easy ERP modulation, and updating bias after easy trials.

### Study 2

Study 2 was a replication of Study 1, and the same software and hardware were used for administering this experiment. Fifty-one participants (26 non-autistics and 25 ASD) took part in this replication (40 males and 11 females total) (Table 2). Of these, 18 non-autistics and 18 ASD had previously participated in Study 1. Eight non-autistic and seven ASD participants participated in Study 2 but not in Study 1; these numbers were too small for statistical analyses of a completely independent pool of participants, so all group comparisons for Study 2 include both repeat and new participants. The time interval between Study 1 and Study 2 was 1-2 years.

**Table 2:** Demographic information for the participants of the Study 2. For age, Block Design, and AQ50 the mean is specified for each group, with the standard deviation in parentheses. The age range is also included in brackets for each group. *** p < 0.001 based on a Student’s t-test.

The number of trials used in the analyses was comparable to Study 1. For the feedback-related negativity (FRN) analysis, non-autistics had a median of 96 correct trials (range of 70 - 116) and a median of 25 incorrect trials (14 - 51), while ASD participants had a median of 88 correct trials (60 - 116) and a median of 32 incorrect trials (5 – 54). For the peak-to-peak analysis of the responses to feedback, non-autistics had median of 59 hard trials (47 – 75) and a median of 60 easy trials (45 – 72), while ASD participants had a median of 60 hard trials (48 – 74) and a median of 58 easy trials (44 – 69). For the analysis of behavioral recency bias, non-autistics had a median of 40 trials when the previous trial was difficult and correct (31 – 60) and a median of 52 trials when previously easy and correct (34 – 65). For ASD participants, the median was 38 trials when previously hard and correct (27 – 49) and the median was 48 trials when previously easy and correct (31 – 64).

To assess the consistency of the measures between sessions, we looked at the correlation between the measures to quantify test-retest reliability. Additionally, we looked at the correlation between measures within each group to test if the measures could originate from a common neural process. If measures did correlate, it would imply that they vary together on an individual-subject level, suggesting that a single mechanism might modulate both.

#### Group classification

To explore group-level classification, we trained a linear support vector machine (SVM) using scikit-learn (Python). We chose this model architecture because it performed consistently well across a randomized set of data splits (80% for training, 20% for testing) and performed at least as well as alternative approaches, including binary logistic regression, random forest, decision trees, gradient boosting, and SVMs with second-degree polynomial features. The feature set included behavioral recency bias after correct and easy trials, the ERP peak-to-peak amplitude difference between easy and hard trials, and FRN magnitude. The preprocessing pipeline involved z-scoring based on the training set. The regularization parameter (C), which controls the trade-off between margin size and classification error, was tuned using a grid search over logarithmically spaced values between 0.01 and 100, with 5-fold cross-validation on the training set. Model performance was evaluated using accuracy, as well as balanced accuracy and area under the ROC curve (AUC) on a held-out test set. Statistical significance was assessed with 1000-iteration permutation testing. For the main analysis, we performed 20 stratified random 80/20 train–test splits and reported average performance across splits. We then examined a representative model using the hyperparameter setting with model performance closest to the median performance across splits, and we used this model to assess the contribution of each feature. Additional evaluations tested generalization across participants (training on unique individuals and testing on non-unique, or overlapping, individuals). Feature contribution was assessed by inspecting the weights associated with individual features and calculating the amount of impact each weight had on the data. In general, linear SVM classifiers are trained to choose the weights w associated with each feature so that the decision function best classifies positive values of *f(x)* to one class and negative values to the other, where *x* is a vector for each sample containing feature values and *b* is a (constant) bias term (here, non-autistic and ASD are the positive and negative classes respectively; Bishop, 2006). By isolating just *w*^⊤^*x* for each feature, we can compare the feature’s contribution to the classifier, with standard caveats regarding weight interpretability (Haufe et al., 2014). A feature that is a good discriminator (and has low class overlap) should transform data from a non-autistic participant to a positive value and an ASD participant to a negative value. We plot these feature contributions using a kernel density estimator (KDE; Bishop, 2006) in Figure 6B-D.

## Results

### Event-related potential and behavioral responses to stimuli are similar for people with and without ASD

Participants performed two-tone pitch discrimination, during which we recorded EEG. On each trial, two pure tones with different frequencies were presented serially. Participants were asked to determine which of the two tones had a higher pitch (Figure 1A, in the example the second tone frequency is “higher”). Upon responding, they were immediately presented with visual feedback: “Correct” in green text or “Incorrect” in red text. Since we were interested in the ability to track performance, which might depend on performance level, we selected a task that produces similar success rates for the two groups but strikingly different effects of trial history (Lieder et al., 2019). We used a simple temporal protocol with two-second intervals both between stimuli and between trials to minimize possible effects of within-trial changes in response amplitude that may differ in neurodivergent groups (Jaffe-Dax et al., 2017).

With these intervals, the peaks of the three main EEG response components (N1 between 70-130 ms, P2 between 150-230 ms, P3 between 250-350 ms) to the two tones (1^st^ and 2^nd^) were similar in the two groups (Rank-sum test: 1^st^ N1, z = -0.415, p = 0.678; 2^nd^ N1, z = -1.55, p = 0.122; 1^st^ P2, z = 0.48, p = 0.631; 2^nd^ P2, z = -1.07, p = 0.284). For N1, the magnitude did not significantly differ between stimuli within a trial (signed-rank test: non-autistics, z = -1.80, p = 0.072; ASD, z = 0.24, p = 0.813). For P2, the magnitude following the second tone in a trial was significantly smaller than for the first tone, but only for non-autistics (z = -3.10, p = 0.002) and not ASD (z = -0.40, p = 0.688). This group difference reached significance, but it was a moderate effect (rank-sum: z = -2.11, p = 0.035, d = 0.54). As expected, both groups had a P3 component following the second tone (Figure 1B), which characterizes time points where a categorical decision is made ("which tone was higher?"; Nahum et al., 2010; Cohen et al., 2013; Jaffe-Dax et al., 2017). The P3 magnitude was similar for the two groups (1^st^ P3, z = -1.05, p = 0.295; 2^nd^ P3, z = -0.60, p = 0.549).

Behaviorally, both groups had similar median success rates in the task (Figure 1C; non-autistics = 71.9%, ASD = 70.0%; rank-sum test: z = 0.76, p = 0.447). However, accuracy was more variable in the ASD group than in the NT group (standard deviation: non-autistics = 0.072, ASD = 0.126; permutation test, p = 0.001), consistent with heterogeneous performance within the ASD sample. Several ASD participants performed very well; those with fewer than five errors were excluded from subsequent group analyses because they did not provide enough Incorrect-feedback trials for a reliable FRN estimation.

### The Feedback-Related Negativity (FRN) is significantly weaker in ASD

Under similar task performance and comparable EEG responses to tones in the two groups, we next asked whether the response to external feedback is also similar. The visual feedback was provided immediately (within 20 ms) after the participant’s button press. To quantify the EEG response to the feedback, we aligned the ERP to the participants’ button press. We rejected three ASD participants who performed extremely well in the experiment and thus had too few incorrect answers to get a reliable FRN response (< 5 incorrect trials; see Figure 1C, the three participants who performed better than all others). The EEG response following “incorrect” feedback is known as the “feedback-related negativity” (FRN) (Miltner et al., 1997). The topography of activity at the time of the negativity (shown in the inset in Figure 2B) indicates the involvement of frontocentral cortical regions (Dehaene et al., 1994; Miltner et al., 1997). Follow-up analyses indicated no differences in results when EEG channels neighboring Cz were considered (Supplementary Figure 1). Additionally, cluster-based permutation testing and multivariate pattern analysis revealed significant effects of the frontocentral cluster of channels in highlighting the group difference in FRN magnitude (Supplementary Figures 2 and 3).

**Figure 2:**
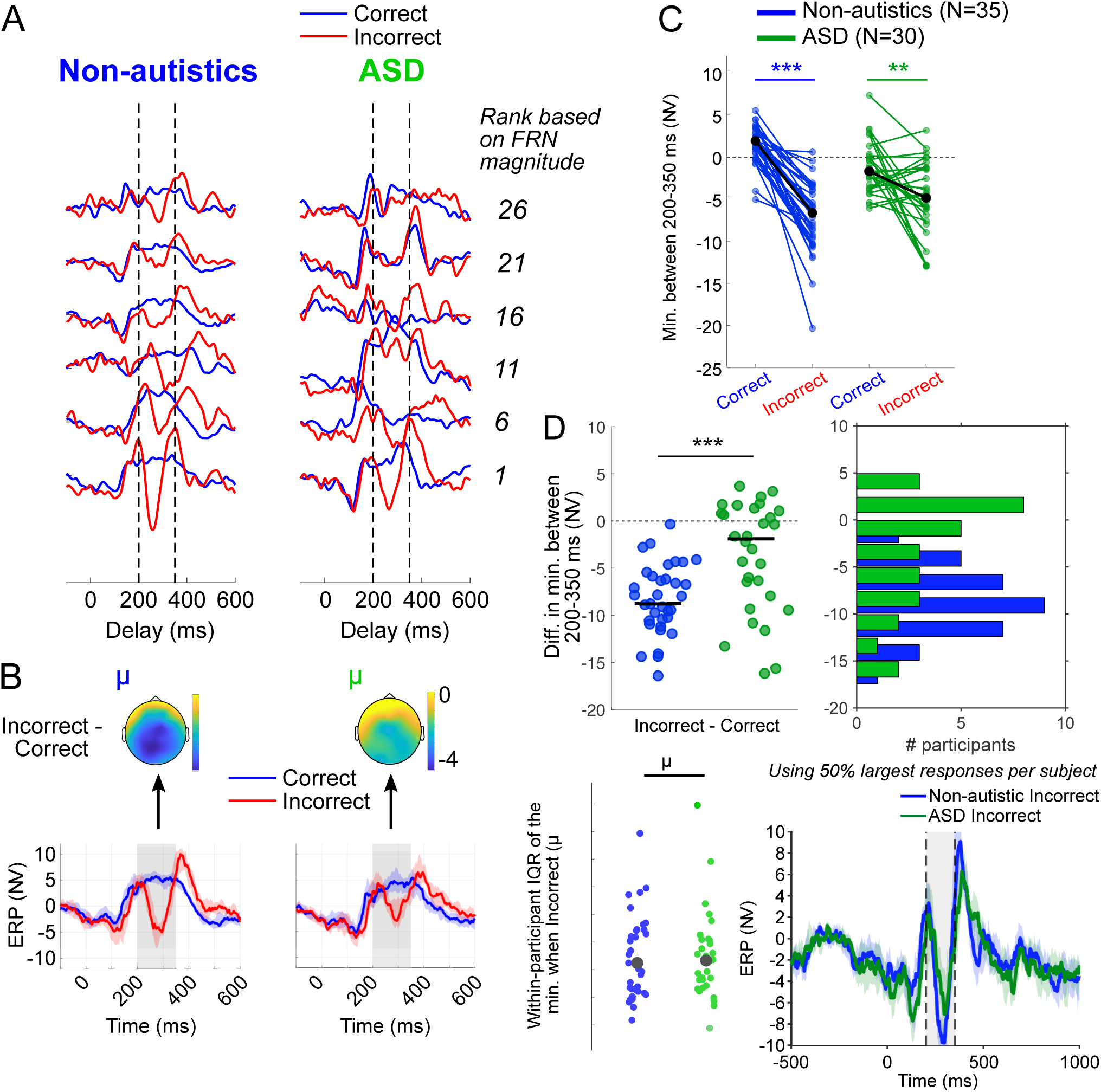
ASD participants’ EEG activity in response to external feedback (Feedback-Related Negativity, FRN) is weaker than that of non-autistics’. (A) Single participants’ FRN, defined as the difference between (minimal) responses following "Correct" (blue) and "Incorrect" (red) feedback within 200-350 ms after their behavioral response (range marked by the dashed lines). FRNs were ranked according to these differences. We plotted responses of six participants from each group, whose ranked magnitude of FRNs were matched with respect to their group (non-autistics on the left, ASD on the right). These ranks are denoted on the right (1^st^ being largest). While all six non-autistics had an FRN, only two of the six plotted autistics’ responses manifest a clear FRN. (B) Median ERP across participants, shaded error bars indicate the bootstrapped 95% confidence intervals of the median. The topography of the difference in voltage between Incorrect and Correct trials averaged within this time interval is shown above each plot. (C) Quantification of the FRN magnitude as the difference between the ERP minimum for incorrect and correct trials, within the range 200-350 ms as indicated in A and B, plotted separately for each participant. At the group level both groups show a significant FRN. (D) Individual dots show the difference in magnitude between correct and incorrect, and the horizontal bars denote the median of each group. The group difference in FRN is highly significant (*** = p < 0.001). To the right is a histogram of these FRN magnitudes, highlighting the difference between groups. (E) Within-subject interquartile range (IQR) of the negativity magnitude between 200 – 350 ms during incorrect trials. Colored dots indicate individual participants, and the black circles indicate the median IQR for each group. (F) For each participant, we calculated the Incorrect response using the half to the trials with the largest negativities (200-350 ms). The negativity is still smaller in ASD than the non-autistics despite analyzing their largest responses.

Figure 2A plots samples of 12 single participants, six with ASD (right) and six non-autistics (left), matched for the rank order of their FRN with respect to their group. It illustrates that nearly all of our non-autistic participants (sampled here by 1^st^ to 26^th^ in descending FRN magnitude) have a clear and large FRN, whereas most ASD (also sampled here by 1^st^ to 26^th^) do not. To quantify the magnitude of the FRN, we determined for each participant the minimum of the response between 200-350 ms after the participant’s button press (Figure 2C). This time window captured the span of the negativity for all participants (dashed vertical lines in Figure 2A, average ERP in Figure 2B). FRN magnitude was then quantified as the difference between the minimum in incorrect and the minimum in correct trials. Figure 2C indicates that in both groups the minimum for the evoked response to incorrect trials was more negative than to positive trials (signed-rank test: Non-autistic, z = 5.16, p < 0.001; ASD, z = 2.62, p = 0.0087). However, the difference was significantly smaller in ASD participants than in non-autistics (Figure 2D, rank-sum test: z = -3.55, p < 0.001). Importantly, the effect size of this group difference was large (Cohen’s d = 0.98). For reference, the effect size (Cohen’s d) of AQ, a questionnaire specifically designed to detect ASD traits (Baron-Cohen et al., 2001), was 1.10 for our participants, comparable to the effect size of the FRN measure. We did not find a correlation within each group between the AQ score and FRN magnitude (Non-autistics, ρ = -0.27, p = 0.13; ASD, ρ = -0.14, p = 0.94); in non-autistics it was shown that “incorrect” feedback yields a larger error- or feedback-related negativity when the task is easier (Miltner et al., 1997) and when errors are more unexpected (Gehring et al., 1993; Holroyd and Coles, 2002), perhaps because unexpected errors are more surprising. This observation suggests that people with ASD may attribute smaller weights to their perceptual predictions and are consequently less surprised (smaller FRN) when they are “surprisingly wrong”.

Another possibility is that the reduced FRN amplitude we observed in ASD is not a neurophysiological characteristic but rather reflects increased variability within and across individuals leading to reduced averages. To test this possibility, we examined both across-participant and within-participant dispersion. No group differences were observed in within-participant variability based on trial-wise FRN minima from Incorrect trials (Mann-Whitney: U = 520, p = 0.953) (Figure 2E). Also, the cross-trial distribution of FRNs could have differed between the two groups; for example, perhaps for ASD participants only a subset of trials had equally large amplitudes to non-autistics, while the FRN was absent in other trials due to a temporary lack of attention. To test whether the group difference reflects a mixture of normal and absent FRN responses in ASD, we repeated the analysis using only the 50% largest (most negative) Incorrect responses for each participant (Figure 2F). The group difference for the overall FRN measure (Incorrect – Correct) remained significant (Mann-Whitney: U = 760, p = 0.002), indicating that it does not reflect a reduced fraction of trials with large FRN responses in ASD. These results indicate that the reduced FRN in ASD reflects a consistent difference in response magnitude rather than increased variability or a subset of absent responses.

Furthermore, we compared quantiles of participants based on their FRN magnitudes. We found that the group difference was not just driven by a subset of the participants; except for the participants with smallest FRN magnitudes, all quantile comparisons exhibited an FRN difference between groups (Supplemental Figure 4). Another account for the group-difference in FRN could be stemming from the coincidentally small number of Incorrect trials, particularly among high performing individuals. However, the FRN group difference remained present event when participants were progressively excluded based on the number of incorrect trials, increasing from a Cohen’s d of 0.90 with all participants (as in our initial analysis) to 1.13 for participants with at least 30 incorrect trials (23 non-autistics, 21 ASD). Also, the magnitude of the FRN was uncorrelated with the number of trials (Spearman’s rank correlation, non-autistics: = 0.107, p = 0.539; ASD: ρ = 0.267, p = 0.153). Thus, the FRN magnitude and the group difference in FRN appeared largely unaffected by the number of trials. Importantly, the this group difference was verified in a follow-up replication study (Study 2) described below.

### Only non-autistics’ EEG responses are modulated by task difficulty

External feedback is often processed in tandem with one’s confidence in their decision (Fleming and Daw, 2017; Grogan et al., 2023). Previous studies of feedback processing have shown that the EEG response can also be modulated by the predictability of making the successful choice, even when the choice produces positive feedback (Holroyd and Coles, 2002; Cohen et al., 2007). Yet in a perceptual discrimination task, unlike past studies focusing on probabilistic rewards, the alignment between one’s internal feedback (confidence) and external feedback is informative regarding the utility of one’s priors. We reasoned that the processing of internal feedback during an EEG response to external feedback might also be altered in ASD. In our task, this would manifest as a difference by task difficulty; if the task in a trial was hard, participants are less confident, and their subsequent response to feedback could be modulated by this confidence.

To test this, we divided the trials into “Hard” and “Easy” (Figure 3A) according to the magnitude of frequency difference in each trial based on the median of the frequency difference in all trials (1.242 semitones was the median frequency difference across all trials and participants). We used the peak-to-peak difference in the evoked response, calculated as the difference between the minimum in 0-150 ms following the participant’s button press and the maximum in 300-450 ms following the button press (see the dashed lines in Figure 3B) to quantify the magnitude of this difference (Figure 3C). The 0-150 ms window was selected to capture the early post-response deflection following the motor decision and feedback-related action, whereas the 300-450 ms window was selected to capture the later positive component of the response-locked ERP. The peak-to-peak measure was therefore used to quantify the overall early-to-late ERP modulation associated with trial difficulty, rather than to isolate a single canonical ERP component.

**Figure 3:**
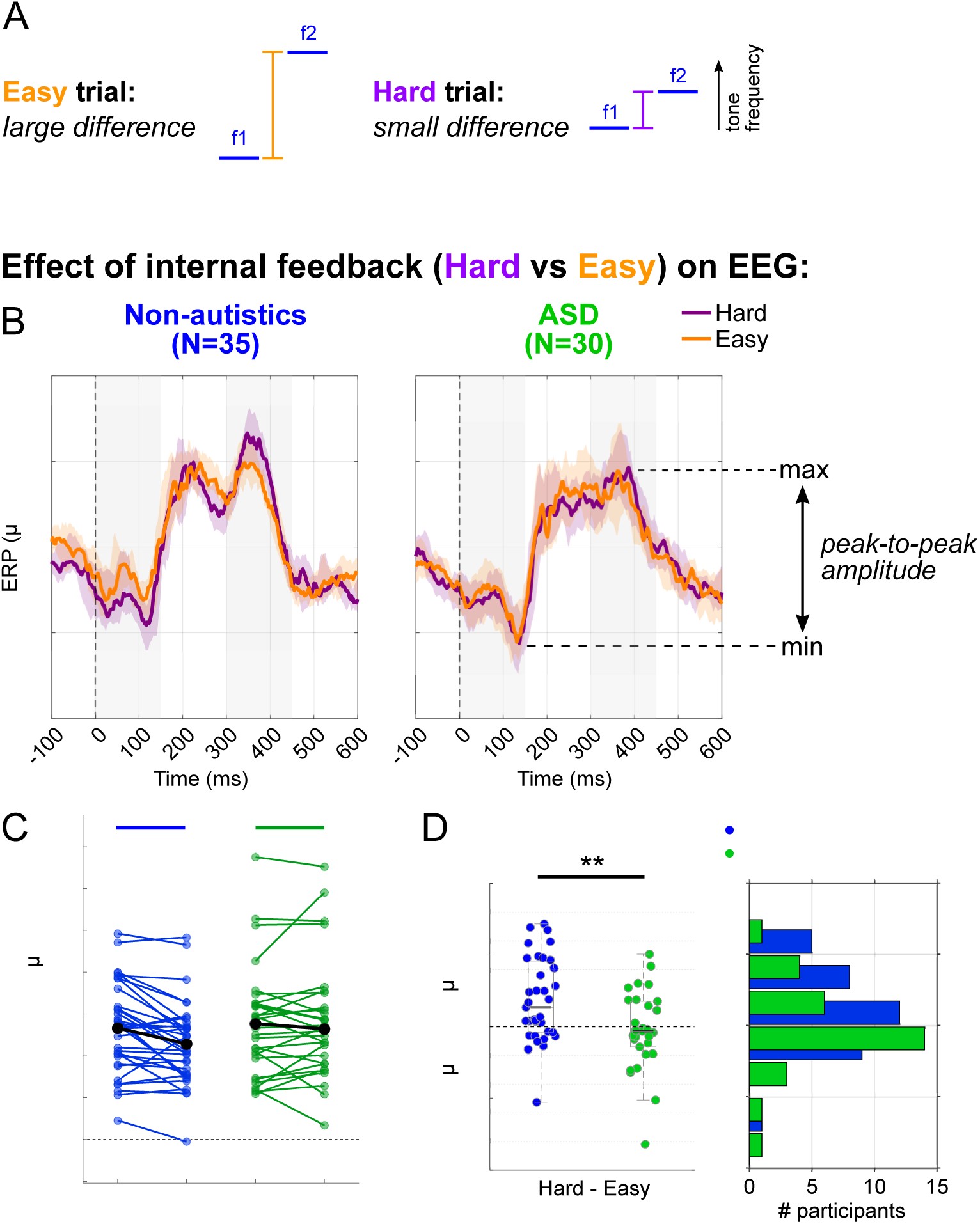
Weaker effects of internal feedback were observed in ASD participants compared to non-autistics. (A) Diagram showing the distinction between easy and hard trials. Easy trials have a larger frequency difference than the median difference across all possible trials (1.242 semitones), while hard trials have frequency differences smaller than this difference. (B) Median evoked responses across participants, time-aligned to the behavioral response, plotted separately for hard (purple) and easy (orange) trials. (C) The peak-to-peak difference in the evoked EEG responses to hard compared to easy trials is significantly larger in non-autistic participants. Note that, here, ASD participants with excellent performance (also excluded from Figure 2) were also rejected to remain consistent with the other measures used later for autism/non-autism classification. (D) Individual dots show the difference in peak-to-peak magnitude between easy and hard trials. On the right is the histogram of these magnitudes (as in Figure 2D). The group difference is significant. ** p < 0.01.

In non-autistics’ EEG responses were modulated by task difficulty (Figure 3B), such that the peak-to-peak amplitude was larger following hard trials compared to easy ones (signed-rank: z = 3.06, p = 0.0022). ASD participants, however, did not show any difference (z = -0.59, p = 0.558). The group difference was significant (rank-sum: z = 2.69, p = 0.0071; Cohen’s d = 0.74). Our results indicate that, in addition to weaker processing of external feedback as evidenced by the FRN, ASD participants also exhibit weaker modulation of the EEG response based on trial difficulty. Taken together, these suggest that the processing of both external and internal feedback are weaker in ASD than in non-autistics.

### Only non-autistics’ perceptual bias is modulated online by task difficulty and success

We examined how external and internal feedback affected participants’ response tendencies in subsequent trials. Previous work had shown that participants’ behavioral responses in a two-interval forced choice task are substantially biased by previous trials’ statistics (Ashourian and Loewenstein, 2011; Raviv et al., 2012, 2014; Akrami et al., 2018; Lieder et al., 2019). Within a Bayesian framework, it was proposed that since the mental representation of the first tone is noisier by the time the 2^nd^ tone is presented, the representation of the 1^st^ tone is contracted more towards an implicit prior (predicted frequency), which is formed by previous trials’ frequencies (Ashourian and Loewenstein, 2011) (though see Boboeva et al., 2024). Because of this, in trials when the first tone is closer to this prior, contraction improves performance (these trials are consequently termed bias+ trials), whereas when it is farther, contraction deteriorates performance (bias-) (Raviv et al., 2012; Akrami et al., 2018) (Figure 4A).

**Figure 4:**
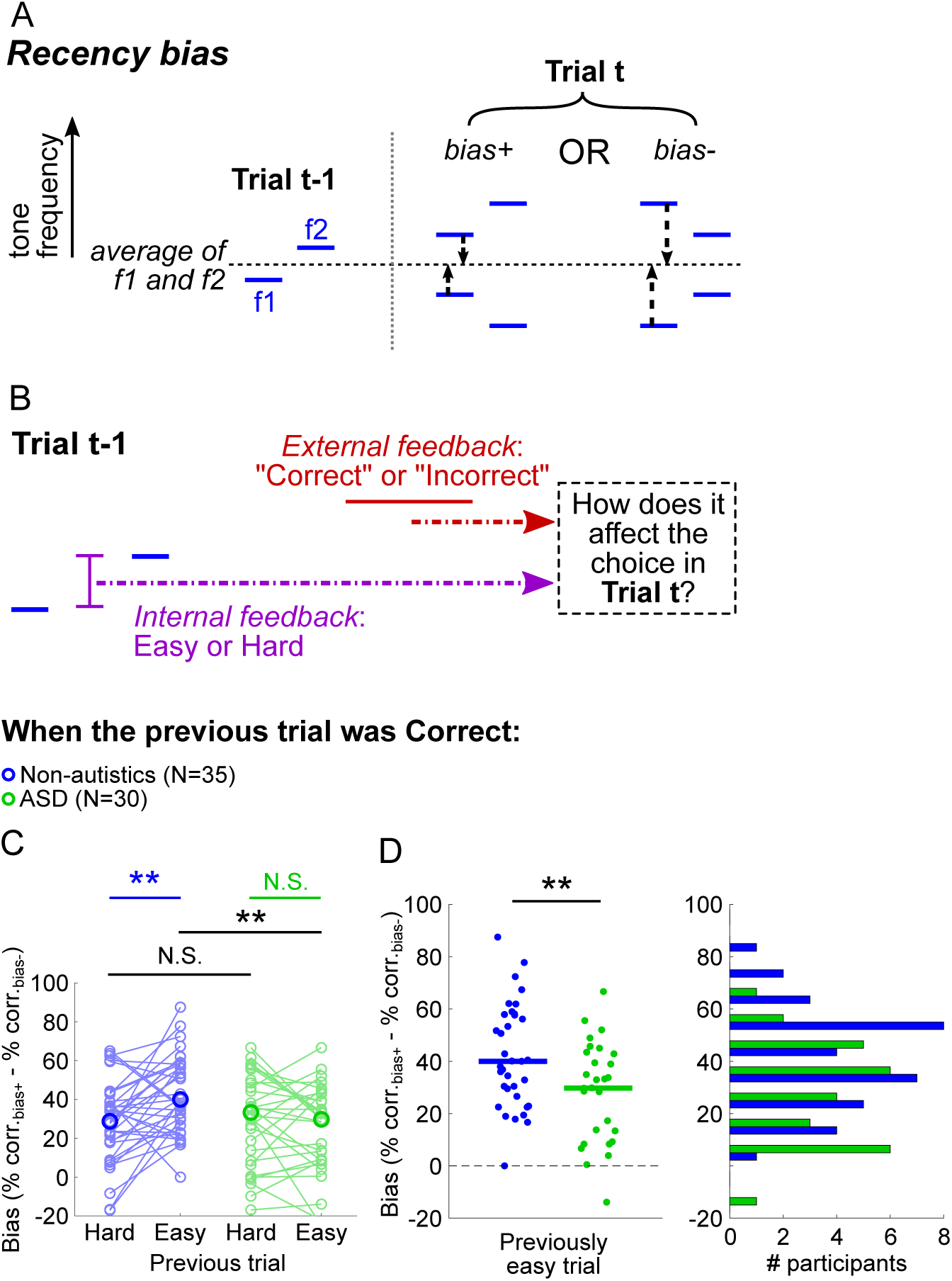
Only non-autistics show a larger recency bias following correct and easy trials. (A) A schematic illustration of recency bias towards recent trial’s frequencies in perceptual decisions. In bias+ trials the first tone’s frequency is closer to the mean frequency of the previous trial, whereas in bias- the second tone is closer. Better performance occurs in “bias+” trials versus “bias-” trials though they differ only in their preceding trial’s frequency. (B) Diagram illustrating the main question of how internal feedback (Hard / Easy) and external feedback (Correct / Incorrect) in the previous trial affect the participant’s choices in the current trial. (C) When non-autistic participants were previously correct yet more confident (easy trials), there was significantly larger recency bias in the subsequent trial. This effect was not found for the ASD group. (D) A closer comparison of the bias difference between groups when the previous trial was both easy and correct, plotted identically to figures 2D and 3D. The group difference was significant. ** p < 0.01

We calculated the recency bias by the difference in success rate between bias+ trials and bias-trials (Ashourian and Loewenstein, 2011; Raviv et al., 2012). In line with previous work (Lieder et al., 2019), the bias for ASD participants was on average smaller than non-autistics’ (rank-sum test: z = 2.39, p = 0.017). Next, we divided these trials based on the difficulty of the previous trial (easy versus hard, based on the threshold of 1.242 semitones) and based on the participant’s accuracy in the previous trial (correct or incorrect) (Figure 4B). All participants were included for previously correct trials. For incorrect trials, we excluded participants who did not have a sufficient number of trials in at least one of the conditions; 32 non-autistics and 28 ASD participants were included (not shown).

We first looked at the effect of the previous trial’s difficulty (internal feedback) on performance, separately for correct and incorrect trials (external feedback). Importantly, we found that non-autistic participants showed a larger recency bias when the previously correct trial was easy compared to when it was hard (signed-rank test: z = - 3.01, p = 0.003). In contrast, ASD participants did not show this effect on bias in correct trials (z = 1.02, p = 0.309), and the bias difference between previously easy and hard trials was significantly weaker than non-autistics’ (z = 2.88, p = 0.004). The recency bias difference was not observed in either group when the previous trial was incorrect (Non-autistics: Incorrect, z = -0.55, p = 0.583; ASD: z = -0.68, p = 0.494). The group difference was particularly driven by a difference in bias when participants were previously correct in an easy trial, where non-autistics showed a larger bias than ASD (z = 2.72, p = 0.006, Cohen’s d = 0.752).

Together, the behavioral group differences suggest that the magnitude of the recency bias and the participant’s subsequent decisions are affected by both the difficulty and accuracy in the previous trial. When non-autistic participants are confident in the stimulus information from the previous trial (after easy) and their confidence has been externally validated (after correct), the effect on the perceptual prior is stronger on their subsequent decisions. ASD participants have weaker processing of internal feedback, and their decisions were unaffected by the accuracy or internal reliability of their response. Importantly, this observation is in line with previous studies reporting reduced online flexibility in updating perceptual priors based on recent events (Sapey-Triomphe et al., 2021). But while studies assessed stimulus statistics, here we assessed the impact of certainty and show that the effect on recency bias is reliable and effective.

### Follow-up study replicates prior results

Study 1 had convincing results for three aspects of online performance monitoring. Given that the two-tone discrimination protocol is composed only of 121 trials, we wanted to verify that these parameters are replicable and that a reliable classification can be done based on these parameters. We therefore invited participants to return for a follow-up assessment with the same protocol approximately 1-2 years after participating in Study 1. Overall, we had 51 participants in this study (26 non-autistics and 25 ASD), and 36 of them were people who also participated in Study 1 (18 NT and 18 ASD), allowing assessment of within participants’ replicability. Four ASD participants in the Study 2 had above 95% trials correct in discrimination – two showed similarly high performance in Study 1, while the other two were newly recruited – and hence were excluded from FRN calculations leaving 18 non-autistics and 16 ASD who participated in both studies. As in Study 1, overall performance was similar for the two groups (mean percent correct in non-autistics = 79%, in ASD = 78%) and correlated across recordings (Non-autistics ρ = 0.5, p = 0.03; ASD ρ = 0.8, p < 0.0001).

All three measures related to feedback and performance monitoring, which were significantly different between groups in Study 1, were also significantly different in Study 2 (Figure 5A-C): FRN magnitude was larger in non-autistics (Figure 5A, z = 2.47, p =0.014, Cohen’s d = 0.741); ASDs’ difference in ERP peak-to-peak magnitude following easy and difficult trials was smaller than for non-autistics (Figure 5B, z = 3.31, p < 0.001, Cohen’s d = 1.044) and ASDs’ reduced flexibility, as manifested in their reduced effect on the recency bias magnitude following easy and correct trials, was smaller (Figure 5C, z = 2.70, p = 0.006, Cohen’s d = 0.883).

**Figure 5:**
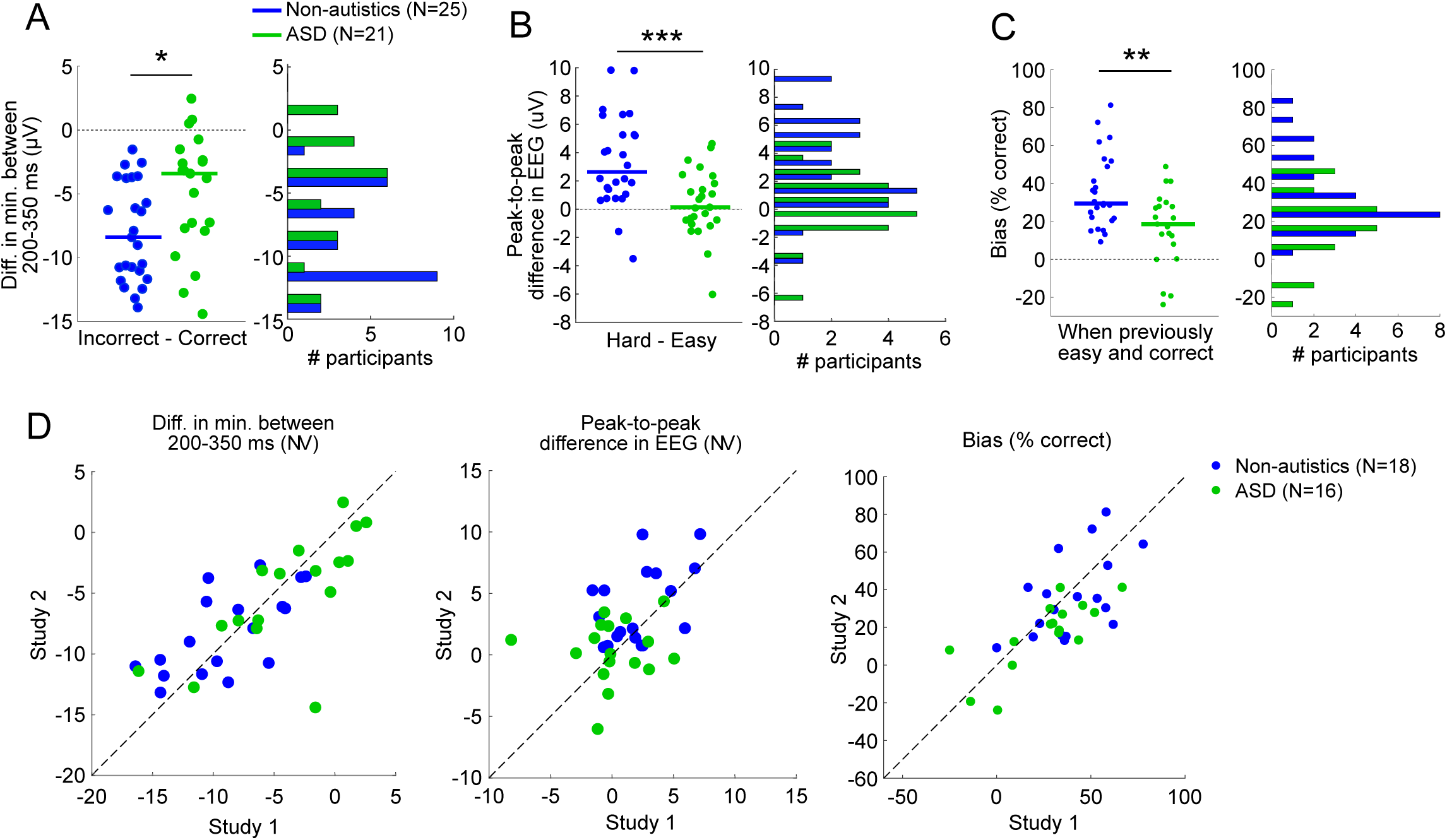
In Study 2, all parameters were replicated and correlated within participants across sessions. (A) Single participant distribution of FRN magnitude, plotted identically to Figure 2D. (B) Single participant distribution of the difference between Hard and Easy peak-to-peak magnitudes of the EEG following the participant’s button press, plotted identically to Figure 3D. (C) Single participants’ distribution of magnitude of online bias updating (following easy and correct trials), plotted identically to Figure 4D. (D) Within participant correlation of these parameters between study 1 and study 2, for the subgroup that participated in both studies. * p < 0.05; ** p < 0.01; *** p < 0.001

Distributional robustness analyses further showed that the group differences were not driven by isolated outliers or a single distributional tail. As we had shown for Study 1, shift-function and difference-asymmetry analyses (Rousselet et al., 2017) showed broadly consistent group differences across the distributions for FRN magnitude, Hard-Easy ERP modulation, and updating bias after Easy trials in both studies (Supplementary Figure 5).

Importantly, for participants who took part in both studies, the key measures showed evidence of test-retest stability more than a year apart; only one measure, the hard-easy peak-to-peak, did not show a significant correlation for ASD participants, but this is expected given that there was on average no peak-to-peak difference within this group (Figure 5D-F; Pearson’s correlations – FRN: Non-autistics, r = 0.656, p = 0.0031; ASD, r = 0.706 p = 0.002; Hard-Easy Peak-to-peak: Non-autistics, r = 0.47, p = 0.049; ASD, r = 0.12, p = 0.66; Bias following easy and correct: Non-autistics, r = 0.540, p = 0.021; ASD, r = 0.767, p < 0.001). Thus, all non-zero measures were replicated at the group level across studies. Individual-level test-retest stability was most consistent for FRN and recency bias.

Given the consistency of these results across replications, we then asked if these measures might be related to a common neural process. To assess this, we combined the participants for study 1 and study 2, taking each participant’s first available session only (42 non-autistics, 34 ASD, after excluding participants with too few incorrect trials), and examined whether the three measures were correlated within each group. For non-autistic participants, the three measures were not significantly correlated with one another (Spearman’s correlations: FRN vs peak-to-peak difference: ρ = -0.078, p = 0.62; FRN vs bias after easy and correct: ρ = 0.06, p = 0.7; peak-to-peak difference vs bias after easy and correct: ρ = 0.11, p = 0.47). In ASD participants, FRN magnitude was weakly associated with both Hard-Easy ERP modulation and recency bias (FRN vs peak-to-peak difference: ρ = -0.32, p = 0.064; FRN vs bias after easy and correct: ρ = -0.36, p = 0.04), whereas Hard-Easy ERP modulation and recency bias were not significantly correlated (ρ = -0.204; p = 0.25). Overall, these mixed findings support the view that the three measures index largely distinct, but possibly not fully independent, aspects of performance monitoring and updating.

### Combining these neurocognitive measures classifies ASD participants

Given that the three performance-monitoring measures were both replicable across studies and captured largely distinct aspects of performance monitoring, we reasoned that a classification model based on all three could be both effective and informative, as it can point to the specific performance monitoring characteristics that are atypical in ASD participants.

We trained a linear support vector machine (SVM) classifier using the three measures as features (for details see Methods). This model choice allows for linear decision boundaries based on the combined contribution of the three features. Over random train-test splits of all participants, the model achieved a mean test accuracy of 78.7% with a mean balanced accuracy of 76.8%. Similarly, the mean receiver operating characteristic (ROC) area under the curve (AUC) was 0.81. AUC represents the probability that a randomly chosen non-autistic participant’s data will be ranked higher by the model than a randomly chosen autistic participant’s data, where an AUC of 0.5 is chance level (Hanley and McNeil, 1982; Figure 6A).

**Figure 6:**
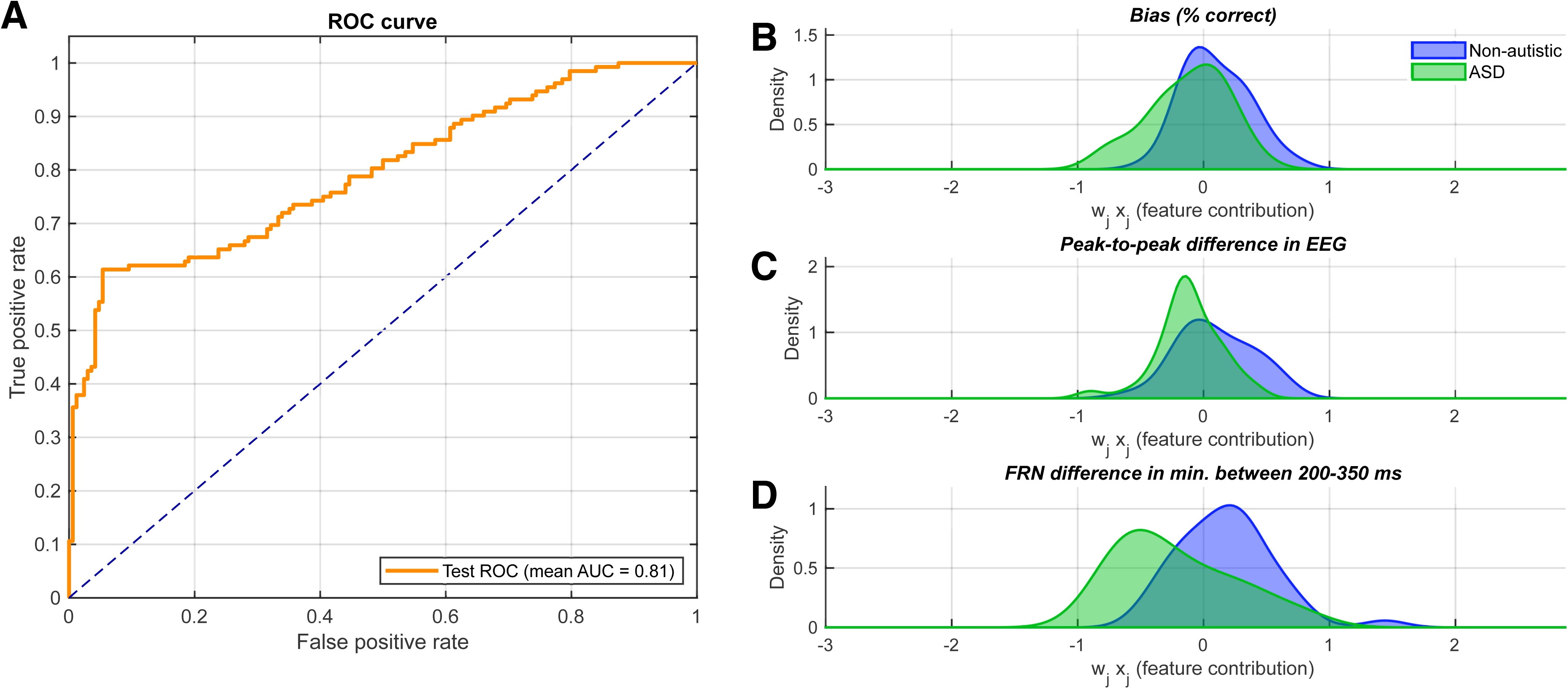
Group classification based on the three parameters of performance monitoring. (A) Receiver operator characteristic (ROC) curve for the representative support vector machine classification model (AUC is 0.81). (B-D) shows the contribution of each main parameter to this classification model, with negative values favoring ASD classification, and positive values favoring non-autistics. The FRN magnitude difference (D) has the largest impact (i.e. largest effect size) since there are fewer areas of overlap between the true classes and each is pushed away, while behavioral bias (B) provides a smaller but still significant contribution. The ERP peak-to-peak amplitude (C) contributes much less.

Next, a representative linear SVM model with performance close to the median accuracy across train-test splits was selected to examine feature contributions to classification. Across train-test splits, this model achieved 78.7% test accuracy and an AUC of 0.81 (Fig. 6B-D; see Methods). FRN and behavioral recency bias following easy and correct trials contributed more to the separation of classes than the peak-to-peak ERP magnitude, where separation can be understood based on the amount of overlap between the density distributions for non-autistic and ASD groups. Performance remained robust when training on participants’ initial experience with the experiment (i.e. all participants in study 1 and non-repeating participants in study 2) and testing on individuals who had already participated once before (participants in study 2 who also completed study 1) with an accuracy of 76.5%, balanced accuracy of 75.3%, and AUC of 0.89. Permutation tests confirmed that classification performance was significantly above chance across all data splits (p < 0.001).

Overall, the results demonstrate that these three behavioral and neurophysiological measures together can dissociate between people with ASD and non-autistics with good accuracy, even for people with mild autistic traits matched for general cognitive scores to non-autistic people.

## Discussion

ASD and non-autistic participants performed the serial pitch-discrimination task with similar overall accuracy and showed comparable sensory-evoked responses to the tones yet differed in how neural and behavioral measures were modulated by trial-by-trial information. ASD participants exhibited a reduced feedback-related negativity (FRN) responses to negative feedback, weaker neural modulation by perceptual difficulty, and diminished modulation of perceptual bias by recent performance. All three effects were replicated in a follow-up study and were stable within individuals across sessions separated by more than one year. However, the low correlations between measures suggest that they capture dissociable aspects of perceptual-decision evaluation rather than a single common mechanism.

The strongest neural finding was the marked reduction in FRN amplitude in autistic participants. FRN was robust in most non-autistic individuals but attenuated or absent in many autistic individuals. Because FRN is generally interpreted as an evaluative response to negative or worse-than-expected outcomes, rather than a sensory response to the feedback display itself (Gehring and Willoughby, 2002; Holroyd and Coles, 2002; Cohen et al., 2007; Walsh and Anderson, 2012; Sambrook and Goslin, 2015), its reduction is unlikely to be explained solely by reduced visual responsiveness. Instead, it is consistent with explicit feedback carrying less evaluative weight in the autistic group. This could reflect reduced surprise, perhaps because predictions were held with less confidence, consistent with reduced pupil responses to unexpected stimuli in autistic humans and mice (Lawson et al., 2017; Noel et al., 2025). Reduced attention or sensitivity to feedback provides another possibility. More generally, the effect may reflect altered weighting of externally provided feedback relative to internally generated task information. The present protocol cannot distinguish between these accounts, yet autistic participants also showed weaker neural differentiation between easy and difficult trials during feedback processing. This reduced sensitivity to internal estimates regarding trial’s difficulty suggests that ASDs’ FRN reduction does not reflect extended attention to internal estimates.

We also found that feedback-evoked responses were modulated by trial difficulty only in non-autistic participants. For non-autistics, larger feedback-evoked responses following difficult judgments suggest that internal estimates of decision difficulty modulated subsequent outcome evaluation. This aligns with other research on EEG responses to external feedback, where larger responses occur when success is more unexpected (Cohen et al., 2007). The EEG modulation was substantially weaker in the autistic group, indicating that internally generated estimates of decision quality exerted less influence on the neural response to feedback. Its lack of correlation with FRN suggests that reduced use of internal estimates of difficulty and reduced evaluative weighting of external feedback may reflect partially dissociable mechanisms for evaluating recent performance. This distinction is important because objective feedback and subjective difficulty provide different information: one indicates whether the decision was correct, whereas the other relates to the participant’s own estimate of its reliability.

The behavioral findings similarly showed reduced modulation of perceptual bias by recent performance and reliability. In non-autistic participants, perceptual bias increased after trials that were both correct and easy. Thus, participants did not simply update their perceptual expectations from recent sensory history; they adjusted the magnitude of its influence according to the inferred reliability of that evidence. This interpretation is consistent with evidence that confidence in recent perceptual decisions modulates serial dependence (Samaha et al., 2019). Easy and correct trials may increase confidence in recent stimulus statistics, thereby increasing their influence on the next judgment. Autistic participants showed substantially weaker modulation. Although they still showed a recency bias, its overall magnitude was smaller and it was not adjusted with increased reliability after easy, correct trials. The group difference therefore concerns both the use of recent history and, more specifically, the adjustment of its influence according to the quality of recent evidence. This behavioral characteristic was reproduced within individuals across sessions separated by more than one year, indicating a stable individual characteristic rather than a transient strategy or measurement noise.

To our knowledge, this specific modulation of perceptual bias by recent trials, whereby its magnitude is enhanced when their reliability is estimated as high, has received little direct attention in autism. It nevertheless connects several previous findings: reduced use of recent information when rapid updating is required (Lieder et al., 2019; Boboeva et al., 2024), atypical responses to volatile environmental statistics (Lawson et al., 2017; Noel et al., 2025), and reduced online correction in motor tasks (Noel et al., 2021; Vishne et al., 2021; Kasten et al., 2023). These effects may therefore involve not only slower updating of priors, but also weaker adjustment by the reliability assigned to recent information based on ongoing performance.

FRN and related performance-monitoring signals have been associated with mediofrontal cortex, particularly anterior cingulate cortex (ACC) (Falkenstein et al., 1991; Gehring et al., 1993; Dehaene et al., 1994; Carter et al., 1998; Botvinick et al., 1999, 2004; Debener et al., 2005). However, our limited EEG measures do not permit source localization. Moreover, the low correlations among measures do not support a common anatomical substrate and may instead indicate contributions from different mediofrontal circuits. Mediofrontal regions remain plausible contributors because of their roles in feedback processing, integration of internal and external information, prediction-error evaluation, and adaptive control (Carter et al., 1998; Botvinick et al., 2001, 2004; Behrens et al., 2007; Beckmann et al., 2009; Oesch et al., 2026). Indeed, structural and functional alterations in these regions have been reported in autism (Thakkar et al., 2008; Simms et al., 2009; Cauda et al., 2011; Chan et al., 2011; Postema et al., 2019; Sapey-Triomphe et al., 2021, 2023). Source-resolved imaging will be needed to identify the regions contributing to each effect.

Previous studies of FRN in autism have yielded mixed results (Hüpen et al., 2016). One potentially important difference between the studies is the relevance of the feedback. Many previous paradigms used monetary rewards or probabilistic outcomes, in which feedback conveyed value, expectancy, or a learned contingency. The added incentives when a reward is associated with a correct and predictable decision may underlie the difference between studies. In the present task, feedback was immediate and veridical: it directly indicated whether the preceding perceptual judgment was correct or incorrect, but correct responses were not associated with a reward. The current findings therefore show reduced neural differentiation between correct and incorrect performance feedback when overall perceptual accuracy is comparable between groups and only task-relevant perceptual information is attached.

An important additional observation is that the three measures which separated ASD and non-autistic participants were not highly correlated in contrast to test re-test within participants. Their low intercorrelations, and complementary contributions to classification, indicate that the group differences were not redundant; each measure captured information not fully contained in the others or in overall task accuracy. However, these results do not establish diagnostic utility, particularly since no other neurodevelopmental or neuropsychiatric populations were examined.

Taken together, these findings suggest that autism is associated with several atypicalities in processes of performance monitoring, where which recent outcomes and estimates of difficulty influence subsequent evaluation and behavior. The stability and relative independence of these effects indicate that they reflect complementary characteristics rather than a single underlying deficit.

## Acknowledgements

This work was funded by the Israel Science Foundation (grant 1731/24) and by the European Research Council (grant 833694). NZ was also funded by the ELSC-SWC Postdoctoral Fellowship, co-advised by MA and AA. YW was also funded by the ELSC PhD Fellowship. KD was funded by the Gatsby Charitable Foundation (GAT3850). AA was funded by the Gatsby Charitable Foundation (GAT3755) and the Wellcome Trust (219627/Z/19/Z).

## Author contributions

NZ, AA, and MA designed the experiment. NZ, JS, YW, VS collected the data. NZ, JS, and KD wrote the analysis scripts and analyzed the data. NZ, JS, KD, AA, and MA drafted the manuscript, and all authors contributed to editing the manuscript.

## Data availability

All EEG and behavioral data from the experiment are available here: https://osf.io/wjbc3. The code used to run the experiment and analyze the data can be found here: https://gitlab.com/auditory-predictive-coding/erp-adaptation. The code used to run the classification analysis can be found here: https://github.com/kiradust/asd-cat.

## Declaration of interests

The authors report no competing interests.

## Notes

### Competing Interest Statement

The authors have declared no competing interest.

### Summary of Updates

Title updated, and reduced biomarker claim throughout. Now use the Huber M-estimator throughout to compute EEG averages. Remove the mini experiment section from the manuscript.

