## Supplementary Information for "Neurocognitive characteristics of perceptual feedback processing differ in autism"

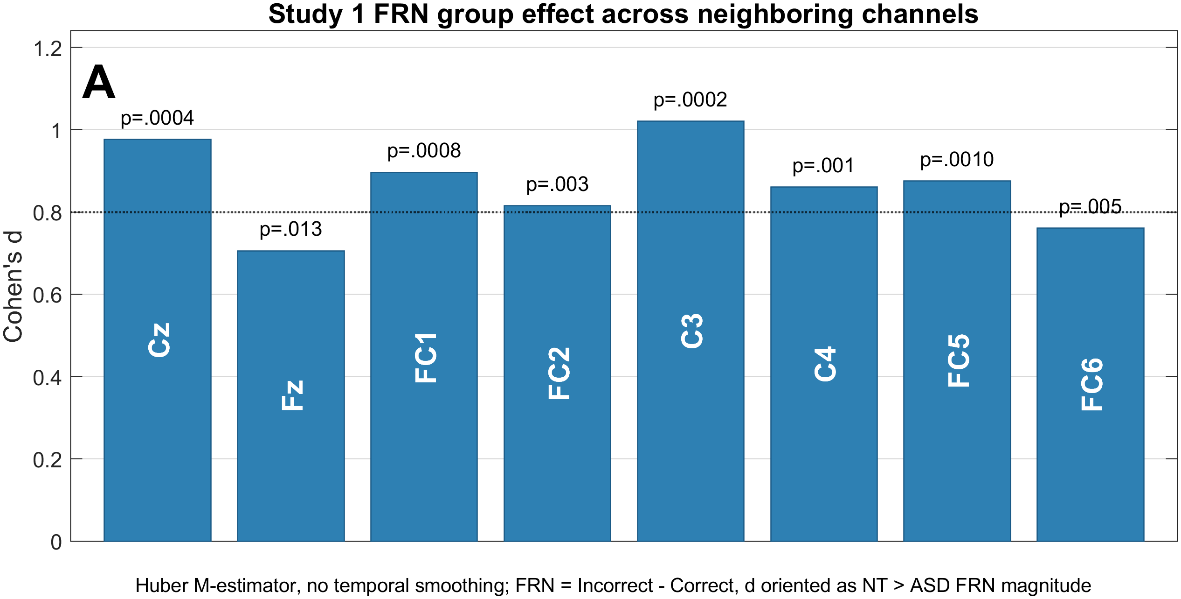

**Supplementary Figure 1: FRN group effect across neighboring channels in Study 1.** The non-autistics, or neurotypicals (NT) versus ASD FRN comparison was repeated at Cz and neighboring fronto-central/central electrodes using the Huber M-estimator without temporal smoothing. Bars show Cohen’s d for FRN magnitude; labels indicate Wilcoxon rank-sum p-values. The effect was significant across neighboring channels and survived FDR correction.

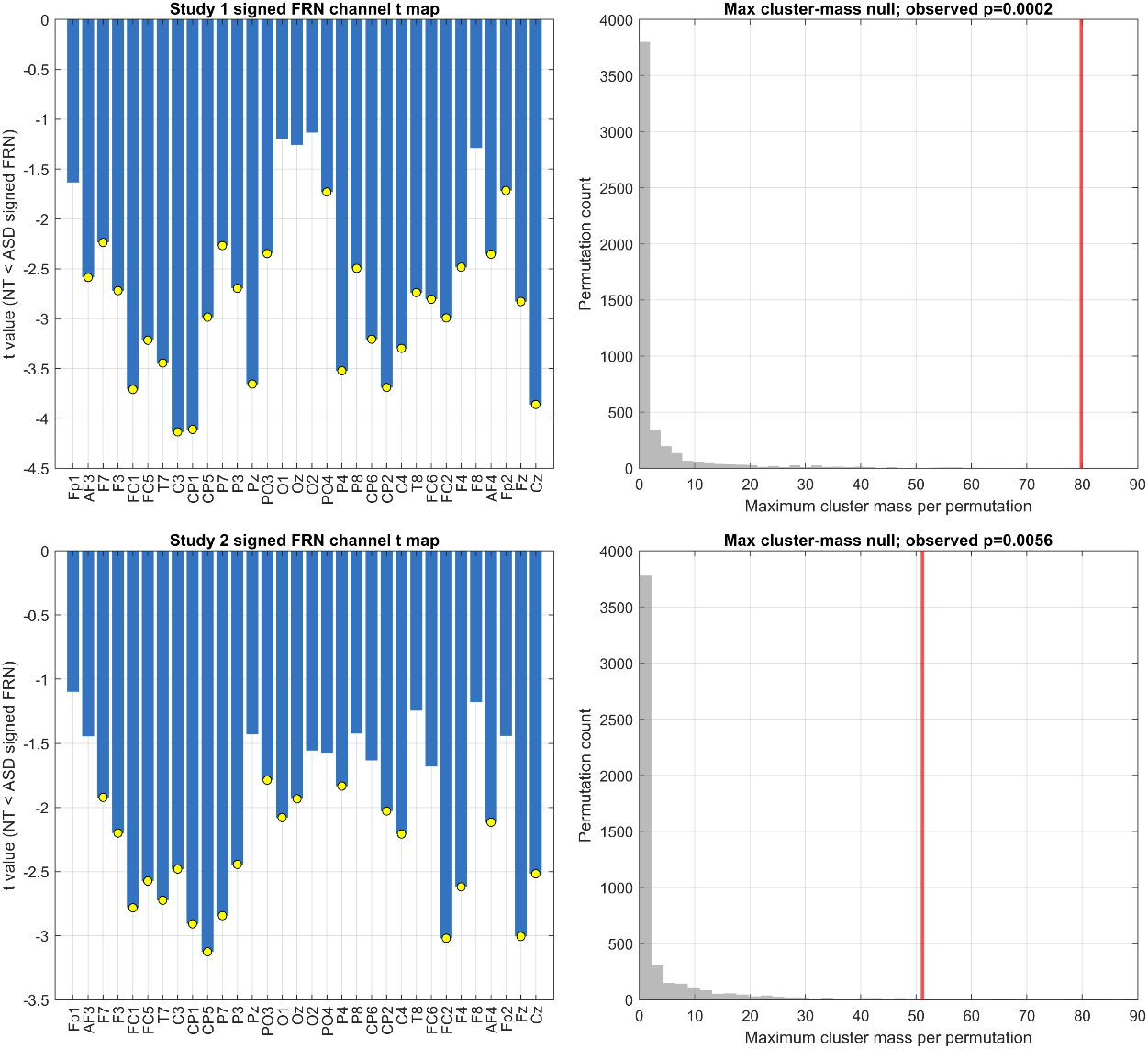

**Supplementary Figure 2: Cluster-based permutation analysis across channels for Study 1.** Spatial cluster-based permutation testing was applied to the NT versus ASD FRN difference across scalp channels. The observed fronto-central cluster, labeled with yellow circles (see left), was tested against a permutation-derived null distribution of maximum cluster mass (right plot). The significant cluster-corrected result indicates that the FRN group difference was spatially contiguous and not restricted to Cz.

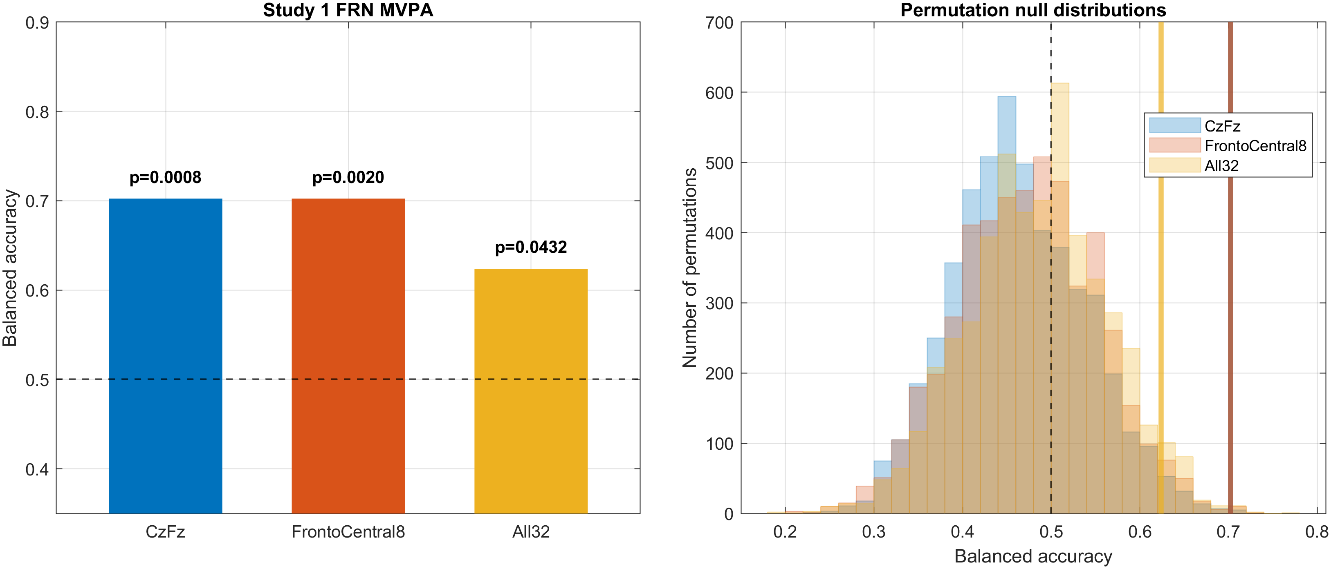

**Supplementary Figure 3: Multichannel multivariate pattern analysis (MVPA) classification of NT versus ASD in Study 1.** MVPA was performed using channel-wise FRN values computed with the Huber M-estimator without temporal smoothing. Linear logistic classification was evaluated using 5-fold cross-validation and label-permutation testing. Left: cross-validated balanced accuracy for NT/ASD classification using Cz/Fz, eight fronto-central channels, or all scalp channels; labels indicate permutation p-values. Right: permutation null distributions of balanced accuracy; vertical solid lines indicate observed balanced accuracy, and the dashed line indicates chance level. The Cz/Fz and fronto-central observed values overlap closely and are therefore partly superimposed. Above-chance classification indicates that the FRN response contained distributed multichannel information distinguishing NT and ASD participants.

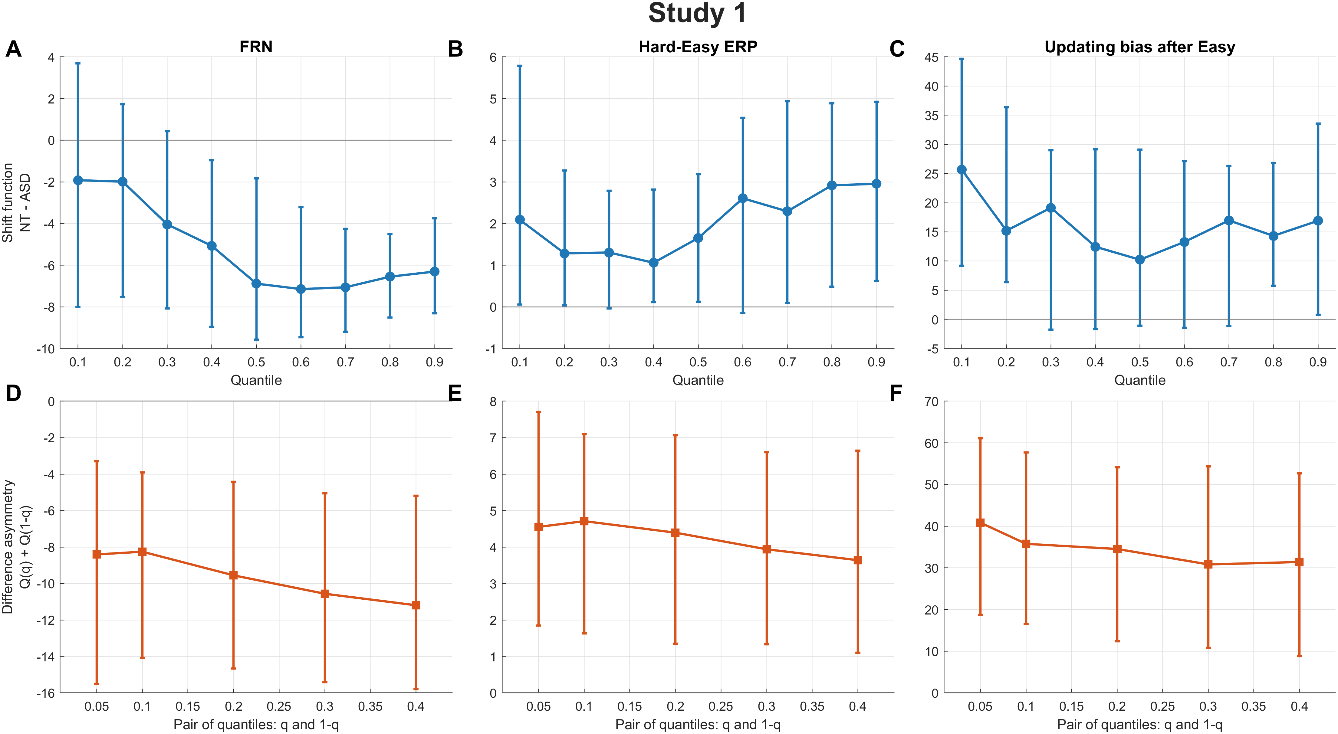

**Supplementary Figure 4: Shift-function and difference-asymmetry analyses for the three primary effects in Study 1.** Shift-function and difference-asymmetry analyses (Rousselet, Pernet, & Wilcox, 2017) were used to test whether the NT versus ASD differences reflected broad distributional shifts rather than extreme participants or a single distributional tail. Top row: shift functions showing quantile-wise NT minus ASD differences with bootstrap 95% confidence intervals. Bottom row: difference-asymmetry functions showing sums of symmetric quantiles of pairwise NT-ASD differences. Columns show 3 measures: FRN, Hard-Easy ERP, and Recency bias (updating bias after Easy Correct trials) – all effects are broadly distributed and not driven by extreme observations or a single distributional tail.

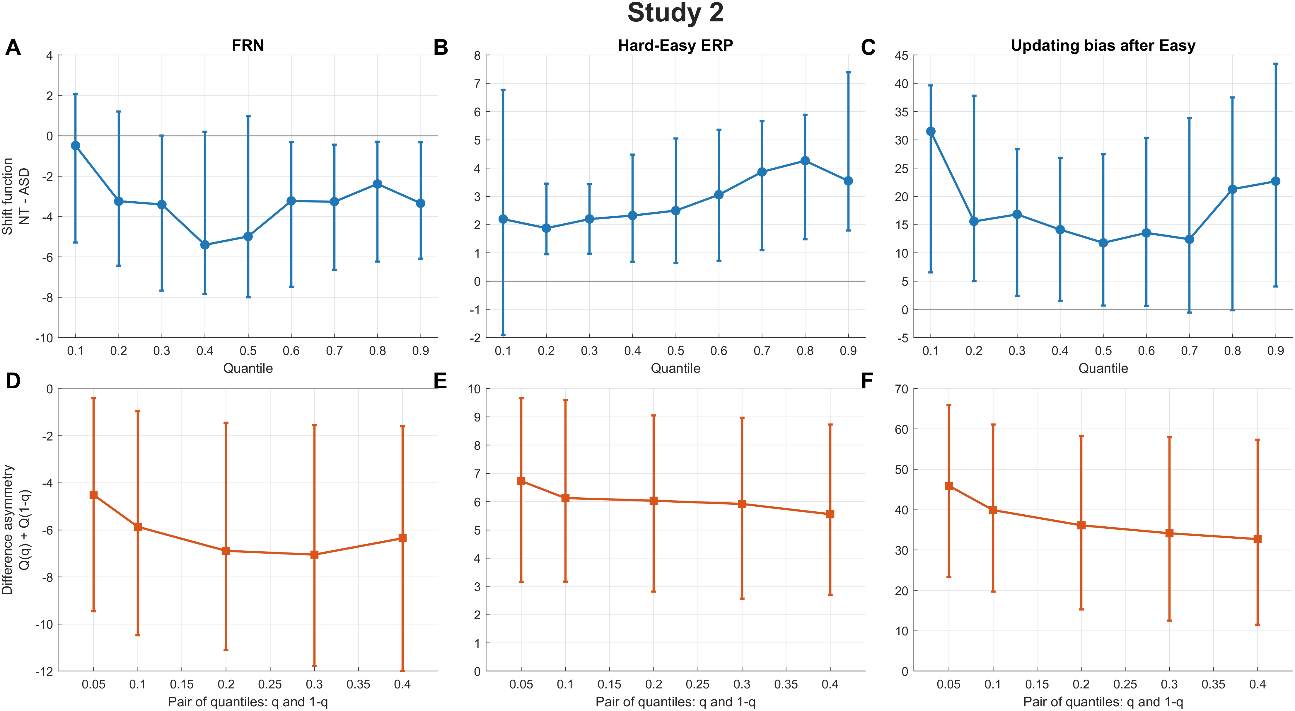

**Supplementary Figure 5: Shift-function and difference-asymmetry analyses for the effects in Study 2**, plotted identically to Supplementary Figure 1.
